# The INO80-EEN complex prevents genomic rearrangements at protein coding genes regions

**DOI:** 10.64898/2026.03.02.709068

**Authors:** M. Bruggeman, S. O. Sall, A. Alioua, S. Graindorge, S. Staerck, J. Muterrer, G. Dupouy, S. Noir, W-H. Shen, J. Molinier

## Abstract

Plants are continuously exposed to a myriad of DNA-damaging agents, including environmental cues such as sunlight. At the cellular level, plants respond to DNA damage by activating DNA damage response (DDR) pathways, in which chromatin remodelers play an important role. Among them, the evolutionary conserved INO80 complex (INO80c) has been shown in Arabidopsis to play a key role in DDR, notably by positively regulating Homologous Recombination (HR). Arabidopsis EIN6 ENHANCER (EEN) is the homolog of Yeast INO EIGHTY SUBUNIT 6 and interacts with the N-terminal region of INO80 in the INO80c. Using plant phenotyping, cellular and molecular biology, and third-generation sequencing technology we investigated how INO80 and EEN regulate plant development and genome integrity. We uncovered new roles for INO80 and EEN in plant growth and for INO80 in fine tuning endoreduplication. In addition, linear genome analysis revealed an important and unexpected function for the INO80-EEN complex in preventing Protein Coding Genes (PCGs) from structural rearrangements in somatic tissue and upon exposure to UV-B. Therefore, our results shed new light on the previously overlooked roles of INO80 and EEN in protecting genome integrity at PCGs.

## INTRODUCTION

Genome integrity is jeopardized in cells by both endogenous and exogenous genotoxic stress. Besides reactive oxygen species metabolic process, endogenous sources are mostly linked to DNA metabolism, either during replication or repair^1^ while exogenous sources come mainly from environmental cues (*e.g.*, sunlight). Due to their sessile life style, plants have to cope with these various sources of DNA damage, with an increasing risk to alter genome integrity. In Arabidopsis, DNA mutation rate in somatic cells is higher than in germline^2^ suggesting the existence of sophisticated mechanisms to signal and repair DNA damages in an error free manner. In eukaryotes, the DNA Damage Response (DDR) signaling relies among other players, on two phosphatidyl-inositol 3-kinase-like (PI3) protein kinases: ATM (Ataxia-telangiectasia-mutated) and ATR (Ataxia-telangiectasia-mutated and Rad3-related^3,4,5^). Defects in DDR and DNA repair pathways lead to enhanced sensitivities to genotoxic stress, resulting in, changes in ploidy levels, nucleotide sequence and linear genome structure, and ultimately leading to cell death^6^.

Chromatin remodelers play important roles in the DDR process^7,8^. Among them, the INO80 complex (INO80c), first identified in a genetic screen in *Saccharomyces cerevisiae* ^9^, is a high molecular weight chromatin remodeler that plays key roles in transcription regulation^10,11^ by the exchange of the histone variant H2A.Z with H2A, affecting nucleosome sliding and the positioning of +1 and -1 nucleosomes at promoters^12,13,14^. The INO80c is now well described with the cryo-EM structure of the core INO80 complex from the fungus *Chaetomium thermophilum* bound to a nucleosome^15^ and with the structural mechanisms for nucleosome binding and sliding^16,17^. Central to this complex is the INO80 ATPase subunit. As in yeast, the Arabidopsis INO80 is involved in transcriptional regulation^18,19^, and in DNA repair by regulating positively Homologous Recombination^20^ (HR). Indeed, INO80 acts in somatic plant cells upon formation of DNA double strand breaks (DSBs) to regulate homologous recombination (HR)^21^. Additionally, INO80 contributes to resistance to DNA damage during replication stress^22^ coordinating cell cycle and DNA repair^23^.

In Arabidopsis, the list of components of the INO80c is increasing with the identification of a dozen of subunits^24^. Among them, the homolog of the yeast INO EIGHTY SUBUNIT 6 (IES6), EEN (EIN6 ENHANCER) was recently identified^25^. IES6 was first studied due to its role in DNA repair^26^. In Arabidopsis, EEN has also been shown to be a subunit of INO80c^24^, controlling chromatin landscape at *EIN2*, a master regulator of the ethylene signaling pathway^27^. However, it remains to be determined whether EEN is associated with cell cycle and/or DDR regulation.

Previous works have convincingly shown the conspicuous role of INO80 in DDR, using genotoxic stress sensitivity assay and transgenic plants carrying reporter constructs or extrachromosomal substrates to monitor error free or error prone DSB repair processes^20,21^. Such approaches did not allow to investigate the DNA repair mechanisms dedicated to the processing of particular DNA damage which may have led to underestimate the role of the INO80c in some repair pathways.

The development of high-throughput long-read sequencing (*i.e.*, Oxford Nanopore Technologies)^28,29^ allows identifying changes in linear genome structure also called structural variations (SVs). Deletions, duplications, insertions, inversions and inversion-duplication are the predominant SVs that are detected with an unprecedented robustness and resolution^30,31^. SVs represent a high contribution to polymorphic variation in many species including humans^32,33^, Drosophila^34^, fission yeast^35^ or tomato^36^. Importantly SVs have been shown to be associated with human cancer^37^ and Alzheimer disease^38^, important agricultural traits (*e.g.*, fruit size) and plant genome evolution^36,39^. However, the underlying molecular mechanisms and the associated factors remain to be deciphered.

In Arabidopsis, by using high-throughput long-read sequencing, we have determined that ATM and ATR repress the microhomology-mediated end joining mechanism^40^ (MMEJ) repair pathway. In this study, we have combined phenotyping, molecular biology and long-read sequencing to decipher the consequences of the loss of function of the Arabidopsis *INO80* and *EEN* on genome integrity.

Our results highlight the role of INO80 and EEN in plant growth and development through the control of cell cycle, and by affecting endoreduplication. Moreover, we identified that INO80 and EEN repress the formation of Inversion-Duplication at protein coding gene (PCG) loci, under normal growth conditions and also upon UV-B exposure.

## RESULTS & DISCUSSION

### *EEN* loss of function affects plant growth and development through cell cycle control

Previous studies have shown that *ino80* mutant plants are impaired in growth and exhibit severe developmental defects^41^. Given that EEN is a subunit of the *Arabidopsis* INO80c^24,25^ we investigated whether this subunit also contributes to the control of plant growth and development. For this, we have analyzed the *een* and *ino80* single mutant plants as well as *ino80 een* double mutant plants grown under laboratory conditions. We found that *een* plants exhibit delay in growth and smaller rosette area compared to wild-type (WT) plants, these phenotypes are even more pronounced in the *ino80* single mutant and in *ino80 een* double mutant plants (Figures 1A-C), suggesting that both mutations might be epistatic. These observations contrast with Zander *et al*^25^ who reported that *een* plants show no phenotype, and highlight that EEN is involved in plant development. To further investigate the involvement of the INO80-EEN complex in plant growth we measured leaf cell area that can reflect cell cycle behavior^42^. The results showed that *ino80*, *een* and *ino80 een* mutant plants exhibit reduced cell size compared to WT plants (Figures 1B and 1D). Similarly, to rosette area, the reduction of cell size of *een* leaves was less pronounced than those of *ino80* plants suggesting that INO80 acts upstream of EEN (Figures 1B and 1D).

**Figure 1:**
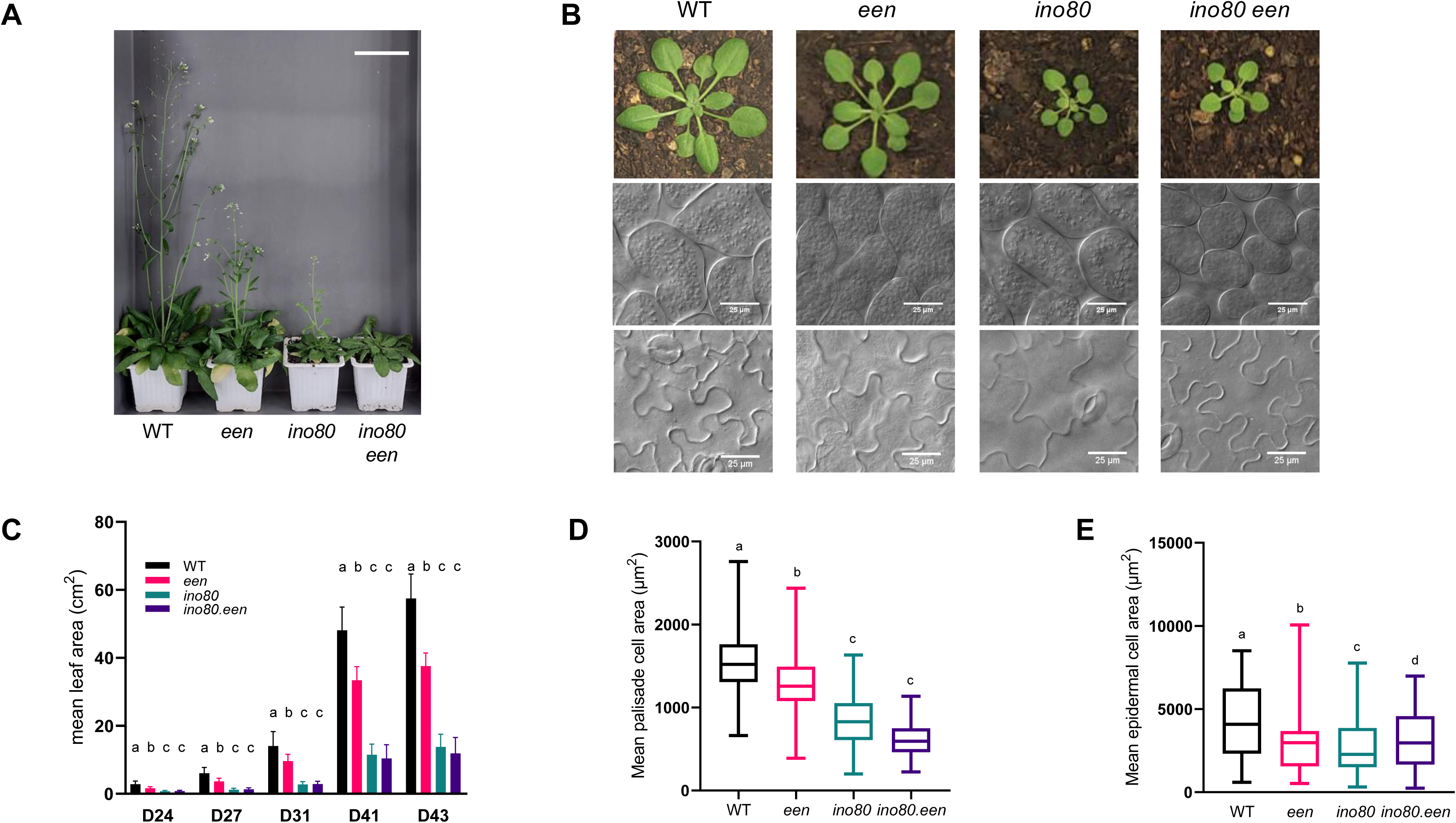
The Arabidopsis *een*, *ino80* and *ino80 een* Arabidopsis mutant plants exhibit alteration in development and cellular phenotypes. A. Representatives pictures of Arabidopsis *een*, *ino80* and *ino80 een* mutant plants compared with wild-type (WT) plants, at 60 days after germination (DAG). Scale bar = 1 cm. **B.** Representative pictures of WT, *een*, *ino80* and *ino80 een* Arabidopsis plants during leaf development at 21 DAG (top panel, scale bar = 7 cm.). Representative microscopy images of palisade (middle panel) and epidermal (bottom panel) cells from the adaxial side of leaves 1 and 2 of WT, *een*, *ino80* and *ino80 een* Arabidopsis plants at 21 DAG. **C.** Histogram representing the mean rosette area (in cm^2^) of WT, *een*, *ino80* and *ino80 een* Arabidopsis plants at different times (D, days) upon germination. Different letters denote significantly different groups (p < 0.05; *One*-way ANOVA with post-hoc Tukey HSD (Honestly Significant Difference) Test). **D.** Box plots representing the palisade cell area (in µm^2^) from leaves 1 and 2 of WT, *een*, *ino80* and *ino80 een* Arabidopsis plants at 21 DAG. Different letters denote significantly different groups (p < 0.05; Mann-Whitney test). **E.** Box plots representing the epidermal cell area (in µm^2^) from leaves 1 and 2 of WT, *een*, *ino80* and *ino80 een* Arabidopsis plants at 21 DAG. Different letters denote significantly different groups (p < 0.05; Mann-Whitney test).

To further characterize the involvement of INO80 and EEN in whole plant development, we performed measurements in root tips that allow monitoring how both cell proliferation and cell expansion are coordinated. Our results show that the root length is affected in the 2 single mutants, as well as, in a similar manner in the double mutant, compared with WT roots (Supplemental Figure 1A-B). Also, both the meristem size and the number of cells within the meristem are significantly reduced in *ino80*, *een* and in *ino80 een* mutant plants compared to WT plants (Supplemental Figures 1C-E) suggesting either a decrease of stem cell niche activity or a faster shift towards elongation /differentiation at the transition zone. Cell area being smaller in *ino80* and *ino80 een* mutants than in the *een* single mutant, it supports the conclusion that INO80 may also act upstream of EEN in root development (Supplemental Figure 1D). These results shed light on the role of *EEN* together with *INO80* in plant growth regulation.

Given that INO80 has been shown to control expression of genes related to cell cycle and regulation of ploidy levels^43,44^, we assumed that EEN could have a similar role. Therefore, we determined whether the expression of key endoreduplication genes were affected in *ino80*, *een* and in *ino80 een* mutant plants. The expression of the transcriptional factor *E2Fc*, that acts as a repressor of cell proliferation and an activator of endoreplication in differentiated cells^45,46^, *KRP6* that regulates CDK activity^47,48^ and *ILP1* that controls ploidy level^49^ were tested by using RT-qPCR. We found they were significantly downregulated in single and double mutant plants compared to WT plants (Supplemental Figures 2A-C). These results are in agreement with the role of INO80 in the transcriptional activation of S-phase genes in *S. pombe*^50^, in cell cycle progression in Arabidopsis^44^ and highlight that EEN also contribute to cell cycle regulation.

Given that increased endoreduplication can be associated to cell size increase in some tissues^51,52,53^, the reduced plant growth and deregulation of cell cycle-related genes identified in *ino80*, *een* and in *ino80 een* mutant plants could be related to impaired ploidy levels. Thus, we used flow cytometry to estimate the DNA content of cells prepared from cotyledons and leaves of *ino80*, *een* and in *ino80 een* mutant plants. We found that *ino80* and *ino80 een* mutant plants exhibit a higher proportion of cells with 2C and 4C DNA contents than WT and *een* plants, in both cotyledons and leaves (Supplemental Figures 2D and 2E). This observation is confirmed by their respective endoreduplication indexes (Supplemental Figure 2F).

**Figure 2:**
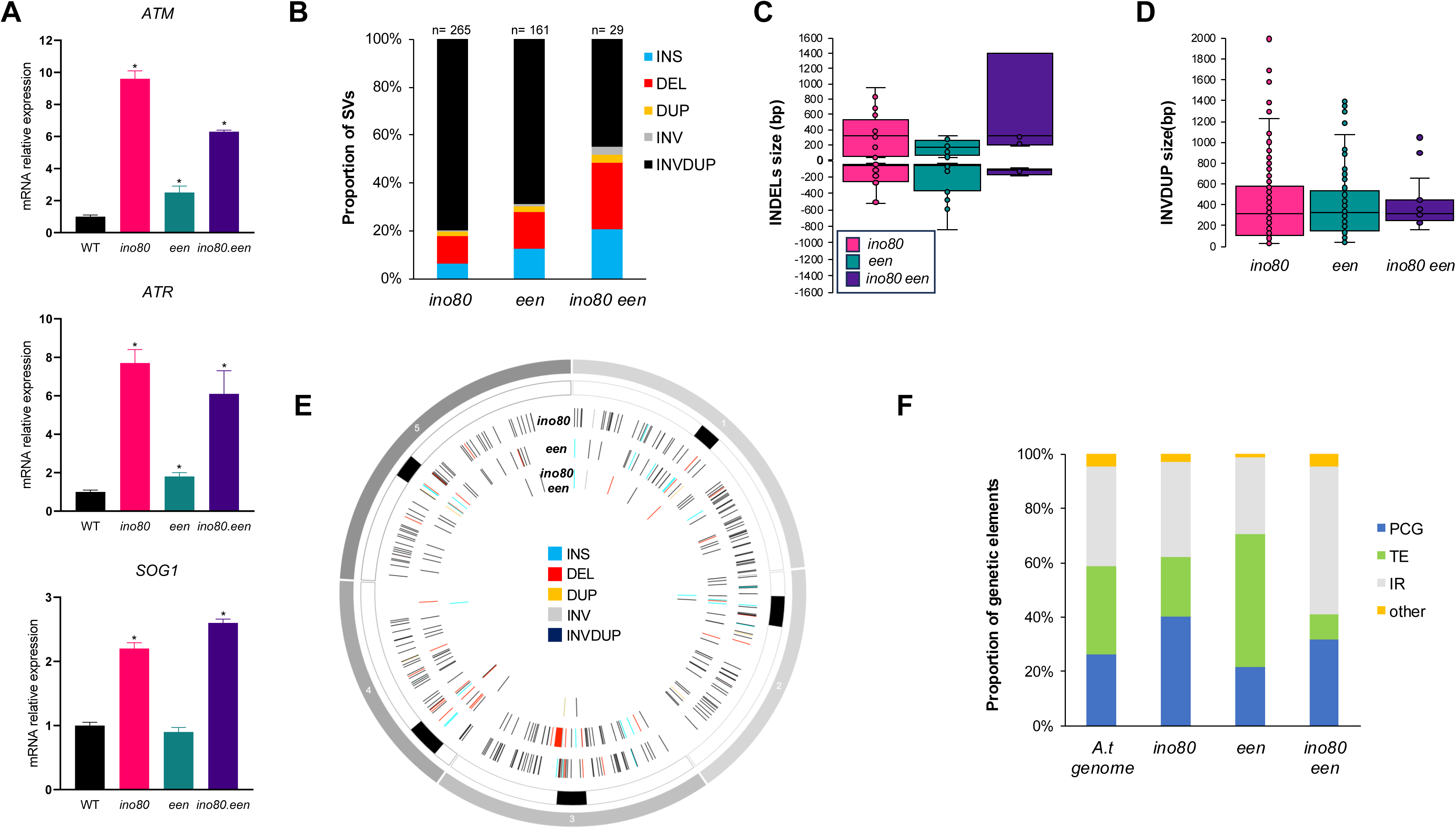
The Arabidopsis *een*, *ino80* and *ino80 een* Arabidopsis mutant plants exhibit constitutive DDR activation and genomic structural variations. A. Histogram representing mRNA steady state level (relative to WT plants) of *ATM*, *ATR* and *SOG1*, in *ino80, een* and *ino80 een* Arabidopsis plants at 21 DAG. (*p < 0.05; t-test). **B.** Histogram representing the distribution of the different types of genomic SV identified in *ino80, een* and *ino80 een* Arabidopsis plants. INS: Insertion; DEL: Deletion; DUP: Duplication; INV: Inversion; INVDUP: Inversion Duplication. n= total number of SV. p= 0.00557 (Chi square test). **C.** Box plots representing the insertions (upper-box) and deletions (lower-box) (INDELs) sizes identified in *ino80, een* and *ino80 een* Arabidopsis plants. p=0.0633 for insertions and p=0.4682 for deletions (*One*-way ANOVA with post-hoc Tukey HSD (Honestly Significant Difference) Test). **D.** same as **C.** for inversion-duplication (INVDUP). p=0.8541 (*One*-way ANOVA with post-hoc Tukey HSD (Honestly Significant Difference) Test) In boxplots, the central line and bounds of the box represent the median and the 25^th^ and 75^th^ quartiles, respectively. The whiskers represent 1.5× interquartile range of the lower or upper quartiles. **E.** Circos representation of genomic SV (INS: Insertion; DEL: Deletion; DUP: Duplication; INV: Inversion; INVDUP: Inversion Duplication) identified in *ino80, een* and *ino80 een* Arabidopsis plants. Black rectangles represent the centromeres. **F.** Histogram representing the distribution of the genetic elements (Protein Coding Genes: PCG; Transposable Elements: TE, Intergenic regions: IR and other elements [*i.e.*, miRNA]) exhibiting SV in in *ino80, een* and *ino80 een* Arabidopsis plants. p= 0 (Chi square test). *A. t* genome represents the distribution of the different genetic elements in the annotated TAIR10 reference genome.

Overall, our results show that both EEN and INO80 are involved in plant growth and cell cycle control, with INO80 playing a particularly important role in the fine-tuning of endoreduplication. Moreover, we found that *een* mutant plants, although exhibiting reduced cell size, show no change in ploidy levels, indicating that distinct cell cycle outcomes occur. These differences may have consequences for genome integrity by affecting the availability of repair templates, such as sister chromatids or homologous chromosomes^69^ reflecting the existence of complex interplays between cell cycle and DNA repair.

### INO80-EEN contributes to genome integrity by preventing the formation of linear genomic rearrangements

INO80 was shown to play an important role in DNA repair in *S. cerevisiae*^10^ and to positively regulate homologous recombination (HR) in Arabidopsis^20^. Next, we aimed to determine whether EEN also acts together with INO80, in pathways involved in the maintenance of genome integrity. First, we determined the expression levels of key factors of the DNA Damage Response (DDR), *ATM* and *ATR* which ensure DDR signaling upon detection of different types of DNA lesions^5^ and *SOG1*a transcriptional factor described as a master regulator of the DDR in plants^68^. A deregulation of these factors would reflect an impairment in DDR signaling with consequences on the DNA repair machinery and likely on the repair outcome^5,54,55^. Upon standard growth conditions, *ATM* and *ATR* mRNA steady state levels are highly enhanced in *ino80* and in *ino80 een* mutants, and more slightly in *een* mutant plants, compared to WT plants (Figure 2A). In the case of *SOG1,* its mRNA steady state level is higher in *ino80* and *ino80 een* mutant plants but not different in *een* mutant, compared to WT plants, (Figure 2A). These results show that *INO80* is a major repressor of the DDR signaling, whereas *EEN* plays a more moderate role.

These data support the essential role of the INO80C in DDR^11^, and highlight a different implication of INO80 and EEN in mediating the DDR signaling. Impairment of the DDR could have major consequences for DNA repair processes by altering the choice of the repair pathways^56,57,58^ that may lead to chromosomal rearrangements.

Secondly, using a third-generation sequencing approach, we generated a genome-wide map of structural variations (SVs) in our mutant lines. Indeed, long reads sequencing technology allows to gain in precision in the linear genome structure and thus to better define the role of INO80 and EEN in maintenance of genome integrity.

Publicly available long reads sequencing data of WT (Col-0; 3 replicates)^40^ in addition to a fourth replicate (this study) have been used as WT-Col-0 reference pedigree to retrieve SVs in our mutant plants. After WT (Col-0) pedigree subtraction, we identified 265, 161 and 29 SVs in *ino80*, *een*, and in *ino80 een* mutant plants, respectively (Figure 2B). Insertions-deletions (INDELS) and Inversion-Duplication (INVDUP) represent at least 90% of the SVs identified in all mutant plants (Figure 2B). While INDELS sizes exhibit significant differences between *ino80*, *een*, and *ino80 een* plants, INVDUP sizes do not display significant size changes (Figure 2C and 2D). In both *ino80* and *een* mutant plants, INVDUP represents more than 60% of the SVs whereas in the double *ino80 een* plants, INVDUP represents around 35% (Figure 2B).

Such difference could be interpreted either as reflecting the control of distinct DNA repair mechanisms by INO80 and EEN, or as resulting from the fact SVs were analyzed only in the 2^nd^ generation of *ino80 een* double mutant plants. In addition, SVs, and more precisely INVDUP, are enriched within chromosome arms (Figure 2E), including PCG, in the *ino80* and *ino80 een* mutants whereas in *een* plants, INVDUP are enriched in TE (Figure 2F). This suggests that upon standard conditions, INO80 rather prevents INVDUP to be formed in PCG whilst EEN would preferentially act at TE. it was reported that INVDUP occurred only at low frequency (< 2%) in WT Arabidopsis plants treated with UV-C and protons as well as in DDR deficient plants (*atm* and *atm atr* mutants)^40^, highlighting a specification in the functions of INO80 and EEN, respectively, to maintain genome surveillance To gain insights into the functions of INO80 and EEN in DDR, we have characterized the *ino80*, *een* and *ino80 een* Arabidopsis plants upon Ultraviolet-B radiation (UV-B) treatment. UV-B as an integral component of sunlight represents a natural source of DNA damage to which plants have naturally developed adapted responses well characterized in the literature^59,60^. In addition, UV-B irradiation stimulates somatic HR in Arabidopsis^61^, suggesting that UV-B can be used as source of DNA damage to further characterize the role of INO80 and EEN in the context of DSB-derived SVs.

We first characterized the phenotypic response of our 3 mutant lines upon UV-B irradiation, by measuring the relative rosette area in comparison with standard growth condition. Conversely to the *uvr3 phrl* double mutant plants, a UV-B hypersensitive mutant, deficient for the expression of the two photolyases involved in the light repair pathway of UV-B-induced DNA damage, the WT plants and the *ino80*, *een* and *ino80 een* mutants did not exhibit a significant difference in leaf area (Figure 3A and Supplemental Figure 3A). In addition, ploidy level was determined and showed that, in all tested genotypes, UV-B exposure did not trigger significant changes (Figure 3B and Supplemental Figure 3B). Thus, this UV-B dose used will be suitable to determine, using long-read sequencing approach, the role of INO80 and EEN in the maintenance of genome integrity upon UV-B exposure.

**Figure 3:**
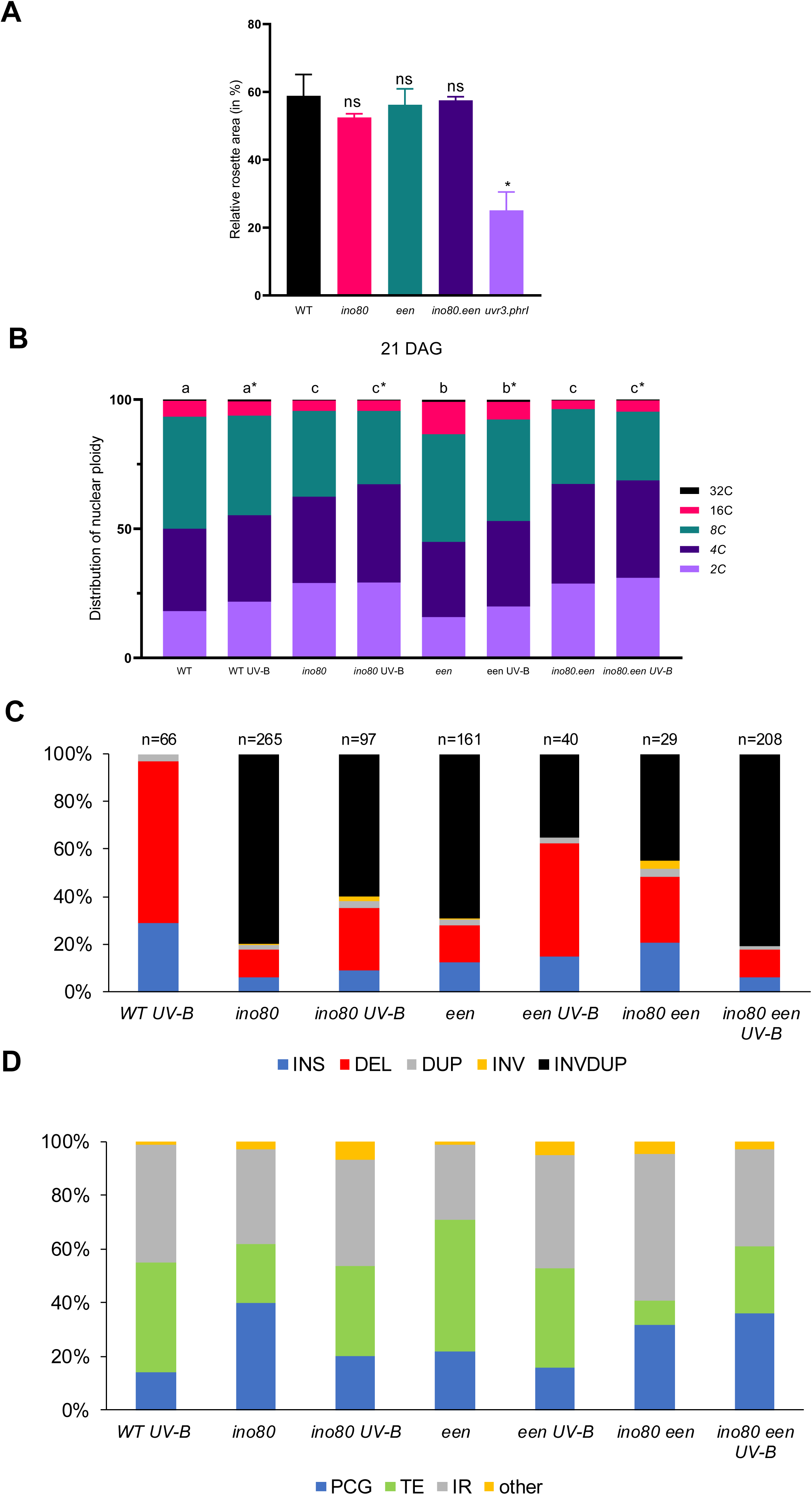
Genomic structural variations and ploidy level in UV-B-treated plants. **A.** Histogram representing the relative rosette area upon UV-B treatment relative to untreated WT, *een*, *ino80* and *ino80 een* plants (*p < 0.05; ns: non-significant; ANOVA with Tukey correction). **B.** Histogram representing the distribution of 2C, 4C, 8C, 16C, or 32C nuclei from leaves 1 and 2 of WT, *een*, *ino80* and *ino80 een* Arabidopsis plants ± UV-B. Different letters denote significantly different groups and asterix (*) indicates significant difference between UV-B-treated samples (Chi square test). **C.** Histogram representing the distribution of the genomic SV identified in WT, *een*, *ino80* and *ino80 een* Arabidopsis plants ± UV-B. INS: Insertion; DEL: Deletion; DUP: Duplication; INV: Inversion; INVDUP: Inversion Duplication. n= total number of SV. **D.** Histogram representing the distribution of the genetic elements (Protein Coding Genes: PCG; Transposable Elements: TE, Intergenic regions: IR and other elements [*i.e.*, miRNA]) exhibiting SV in WT, *een*, *ino80* and *ino80 een* Arabidopsis plants ± UV-B.

After/following WT (Col-0) pedigree subtraction and removal of the SVs identified in each untreated mutant plants, we found 66, 97, 40 and 208 *de novo* SVs in WT-, *ino80-*, *een-* and *ino80 een-*UV-B treated plants, respectively (Figure 3C). UV-B exposure promotes the formation of more than 95% of INDELS in WT plants (Figure 3C), in agreement with the previously published data^40^. The increased frequency of INDELS is also observed in *ino80-* and *een-*UV-B-treated plants albeit INVDUP events still represent 35 and 60%, respectively, of the total SVs (Figure 3C). Upon UV-B exposure whilst deletion size remains unchanged, the insertion sizes significantly vary with a decrease in insertion size in *ino80* and *ino80 een* plants, whereas it causes an increase in *een* plants (Figure 4A). This suggests that different gap filling mechanisms (INO80-dependent and -independent) exist leading to different outcome of repair. Also, conversely to the single mutant plants, the *ino80 een-*UV-B-treated plants exhibit an increase of INVDUP frequency, representing up to 80% of the total number of *de novo* SVs (Figure 3C). While it is observed an opposite occurrence of INVDUP between the 2 single mutants and the double *ino80-een* mutants (Figure 3C), INVDUP size is not affected by UV-B treatment in any of the tested genotypes (Figure 4B). This reflects that INO80 and EEN synergistically prevent the formation INVDUP upon UV-B exposure in *A. thaliana*. UV-B-induced SVs in WT, *ino80* and *een* plants did not show a significant bias within particular genetic elements (PCG, TE or IR; Figure 3D) compared to WT treated plants, whereas in the *ino80 een* double mutant plants SVs remained enriched along chromosome arms and in PCG (Figures 3D and 4C-E).

**Figure 4:**
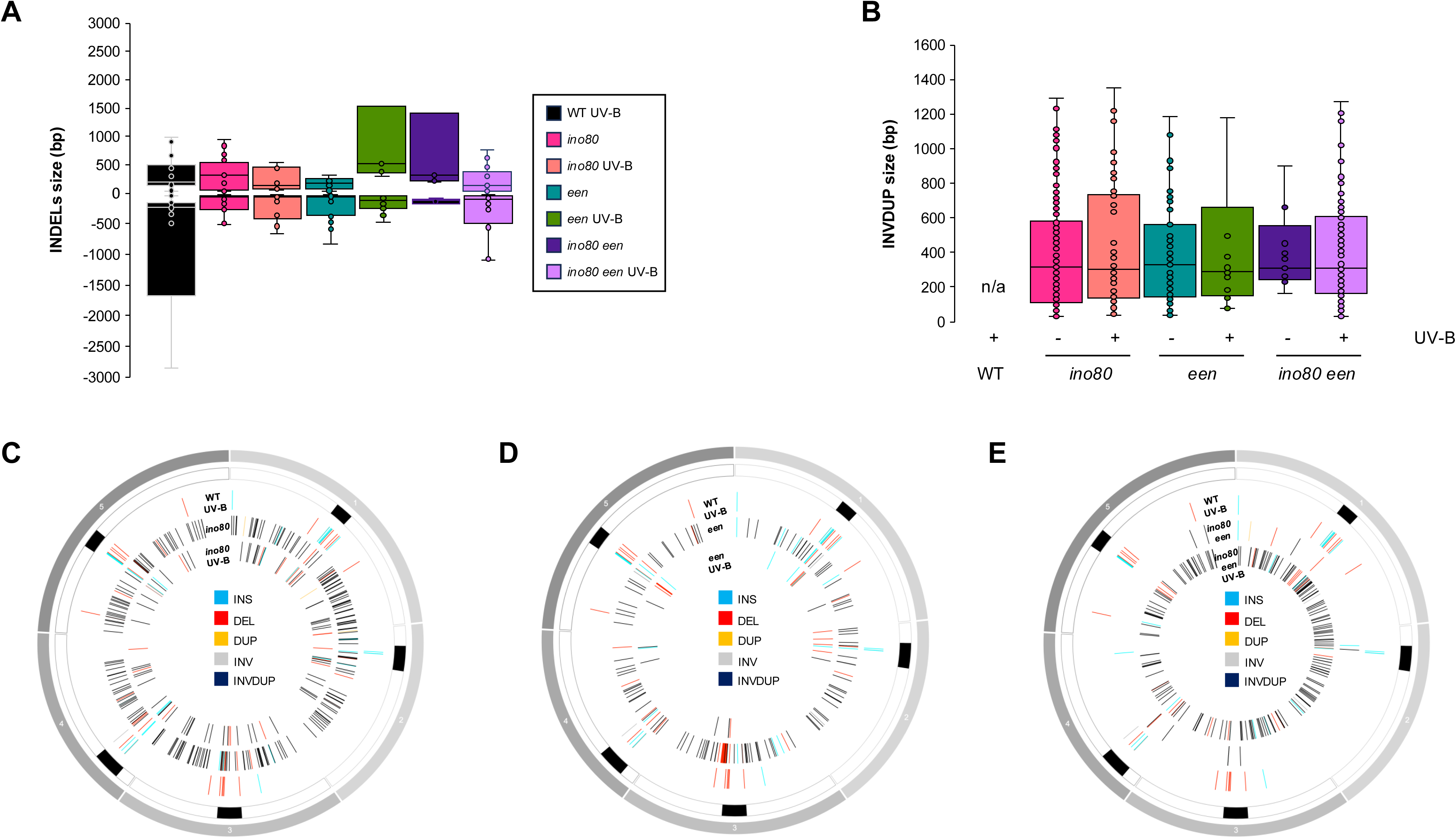
Sizes of genomic structural variations identified in UV-B-treated plants. **A.** Box plots representing the insertions (upper-box) and deletions (lower-box) (INDELs) sizes identified in WT, *een*, *ino80* and *ino80 een* Arabidopsis plants ± UV-B. (p = 0.0140; *One*-way ANOVA with post-hoc Tukey HSD (Honestly Significant Difference) Test). **B.** same as A**D.** for inversion-duplication (INVDUP). n/a: non-applicable. (p = 0.03423; *One*-way ANOVA with post-hoc Tukey HSD (Honestly Significant Difference) Test). In boxplots, the central line and bounds of the box represent the median and the 25^th^ and 75^th^ quartiles, respectively. The whiskers represent 1.5× interquartile range of the lower or upper quartiles. **C.** Circos representation of genomic SV (INS: Insertion; DEL: Deletion; DUP: Duplication; INV: Inversion; INVDUP: Inversion Duplication) identified in WT + UV-B and in *ino80* ± UV-B Arabidopsis plants. **D.** Same as **C.** for WT + UV-B and in *een* ± UV-B Arabidopsis plants. **E.** Same as **C.** for WT + UV-B and in *ino80 een* ± UV-B Arabidopsis plants.

Collectively these results suggest a role for INO80 together with EEN in safeguarding PCG against major structural variations, during plant vegetative growth and upon exposure to UV-B. These results contrast from recent reports that found than more than 80% of the SVs are INDELS and occurred in TE and intergenic regions within constitutive heterochromatin^40^. This holds true in the DDR deficient Arabidopsis plants: *atm*, *atr* and *atm atr*^40^, highlighting that INO80- and EEN-dependent surveillance mechanisms exist to prevent PCG to undergo genomic rearrangements upon UV-B exposure.

Indeed, inverted duplications, also termed foldback inversions, were found to arise from a DSB intermediate and are further categorized in replicative and non-replicative types^62,63,64,65^. It was recently proposed that, in yeast, DSB-induced INVDUP are formed through a DNA polymerase δ-dependent mechanism^66^. Moreover, in Arabidopsis, enhancement of somatic HR frequency has been observed in plants exhibiting reduced expression of the gene coding for the catalytic subunit of the DNA polymerase δ (*POLdelta1*)^67^. Thus, it can be assumed that a mechanism related to foldback inversions exists in *Arabidopsis*, with a prominent, direct or indirect role for INO80 and EEN.

The exact mechanism by which INO80 EEN prevent INVDUP to occur still remain to be deciphered. Nevertheless, these results provide valuable lines of evidence to support the idea that INO80 and EEN play a major role in the surveillance of PCG integrity conversely to ATM and/or ATR^40^.

These findings open new avenues for DNA repair research by identifying novel players in the fold-back repair mechanism. Moreover, they provide a tool for generating genetic diversity at PCGs, particularly for breeding programs.

## MATERIALS & METHODS

### Plant materials and growth conditions

All plants used are in the Columbia ecotype (Col-0). *ino80-5* (SALK_067880) and *een-2* (SALK_129237) plants, referred as *ino80* and *een*, respectively in this study, have been used^25,41^. The *een-2 ino80-5* double mutant plants have been generated by crossing. Homozygotes F3 lines were used for all described experiments. Plants were cultivated *in soil* in a culture chamber under a 16 h light (light intensity ∼150 μmol m^−2^ s^−1^; 21°C) and 8 h dark (19°C) photoperiod. For *in vitro* growth, plants were grown in on solid GM medium [MS salts (Duchefa), 1% sucrose, 0.8% Agar-agar ultrapure (Merck), pH 5.8] under a 16 h light (light intensity ∼80 μmol m^−2^ s^−1^; 21°C) and 8 h dark photoperiod.

### Plant phenotyping

For monitoring root growth, seedlings were germinated and grown *in vitro* on vertical plates using the GM medium. Root length was scored during 5 days and measured using the Fiji software. At least 25 plants per genotype have been used and 2 biological replicates have been performed.

For monitoring of plant rosette leaf area, plants were germinated on soil and transferred to individual pots after 2 weeks. Pictures were taken daily and leaf area was calculated using the Image J plug-in developed by Dr. J. Mutterer: https://github.com/mutterer/services. At least 21 plants per genotype have been used and 2 biological replicates have been performed (data from 1 biological are shown).

### Cytology

Leaves were prepared has described in^70^. Samples were observed using an Axio observer microscope (Zeiss) using DIC optics. Images were captured from at least 6 leaves per genotype. Mean cell area was determined using image J^73^.

Root meristems were mounted in propidium iodide (10 μg/mL) according to the method described in^71^. Image acquisition was performed using the LSM 780 microscope (Zeiss) with a 20X objective. The number of cells in the meristem zone and the length of root meristem were calculated with using Fiji^74^.

### Measurement of ploidy level

The nuclear DNA content was determined from 8-DAG-old cotyledons and leaves 1 and 2 of 21-DAG-old Arabidopsis plants as described in^71^. The Attune Cytometer and the Attune Cytometer software (Life Technologies) was used to record the relative fluorescence intensities of DAPI-stained nuclei. Flow cytometry experiments were performed using 2 independent biological replicates. One biological replicate consists of at least 10 plants grown at the same time period.

### Total RNA extraction and RT-qPCR

Total RNAs were extracted from leaves 1 and 2 of 21-DAG-old plants, using TRI reagent (MRC) following the manufacturer instructions. Reverse transcription reaction was performed using the SuperScript IV Reverse Transcriptase (RT) kit (Thermo Fischer Scientific). qPCR was performed, including technical triplicates, using a Light Cycler 480 and Light Cycler 480 SYBR green Master mix (Eurogentec). Primers are listed in Supplemental Table 1. Experiments were at least duplicated using independent biological replicates (one biological replicate consists of at least 10 plants grown at the same time period and pooled together).

### UV-B irradiation

21-DAG-old Arabidopsis plants (WT, single and double mutant plants) were exposed during 15 min to four bulbs of UV-B Broadband (Philips—TL 40W/12 RS SLV/25) to deliver a total dose of 4500 J/m^2^. Leaves 1 and 2 were harvested 24 h upon irradiation and snap frozen.

### Genomic DNA extraction and library preparation

Genomic DNA was prepared from 100 to 200 mg of leaves using the plant DNA extraction kit Nucleon Phytopure (Cytiva). Ultra-long DNA library for Nanopore sequencing was produced from 100 to 200 fmoles of HMW genomic DNA using the NEBNext companion module (NEB) and the Ligation Native Sequencing Kit V9 (ONT). 5–50 fmoles of the library was loaded onto ONT FLO-MIN R9.4.1 or ONT FLO-PRO R9.4.1 R9.4.1 flow cells.

### Identification of genomic structural variants

Reads were basecalled with ont-guppy-gpu_6.3.8 with the model dna_r9.4.1_450bps_sup.cfg. The analysis was performed using a Snakemake script adapted from the Oxford Nanopore Structural Variant pipeline (https://github.com/nanoporetech/pipeline-structural-variation). Sequencing quality was evaluated with MinIONQC (V 1.33.5). The mapping was performed with Minimap2 (V 2.17) using the A. thaliana, Col0-TAIR10, as reference genome. The mapping quality was checked with Nanoplot (V 1.30.0). Coverage was evaluated by mosdepth (V 0.2.7) and Sniffles (V 1.0.11) was used to identify the structural variations (SV) in comparison with the reference genome Col-0-TAIR10. SVs have been filtered using the following parameters: minimal SV length 1 bp, maximal SV length 1 000 000 bp, minimal read length 1000 bp and minimal mapping quality 20. SVs of the same type (insertion, deletion, duplication, inversion or inversion duplication) with the same genomic coordinates (Chr, start-end) ± 50 bp have been considered as identical. Sequencing statistics are shown in Supplemental Table 2.

## ACKNOWLEDGEMENTS

This research was funded by the CNRS Mission for cross-cutting and interdisciplinary initiatives (MITI) in the frame of the program: Adaptation of the living to its environment program (AAP 2020), by the French National Research Agency (ANR-20-CE20-0021 and ANR-19-CE20-0018), by the EPIPLANT Groupement de Recherche (CNRS, France) and by the International Atomic Energy Agency Coordinated Research Project (D24015): Radiation-induced Crop Diversity and Genetic Associations for Accelerating Variety Development.

Salimata Ousmane Sall received a CNRS PhD grant (Doctorants CNRS 2020 Actions transverses en appui aux défis sociétaux).

## DATA AVAILABILITY

The fast5 files will be available on demand.

## EXPANDED VIEW FIGURE LEGENDS

**Supplemental figure 1:**
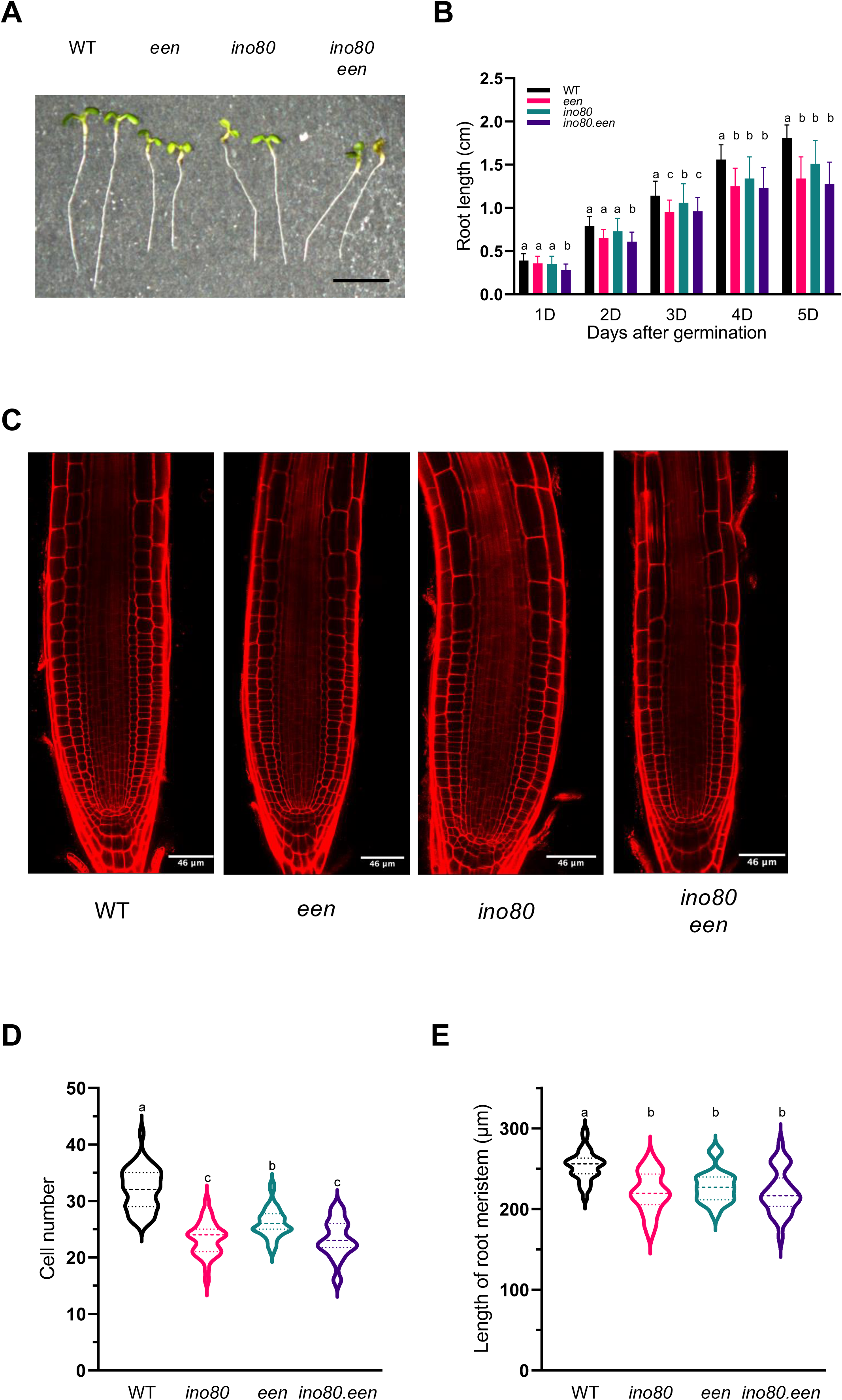
Arabidopsis *ino80, een*, and *ino80 een* plants are affected in their root development. **A.** Representative picture of WT, *ino80, een*, and *ino80 een* Arabidopsis plants seedlings grown vertically at 5 days after germination (DAG). Scale bar = 1 cm **B.** Histogram representing the root length (in cm) of WT, *ino80, een*, and *ino80 een* Arabidopsis plants at 1, 2, 3, 4 and 5 DAG. Different letters denote significantly different groups (p < 0.05, *One*-way ANOVA with post-hoc Tukey HSD (Honestly Significant Difference) Test). **C.** Representative confocal images of propidium iodide-stained roots tips of WT, *ino80, een*, and *ino80 een* Arabidopsis plants. Seedlings were imaged at 5 DAG. Asterix (*) indicates the shift between the meristematic cells and the transition zone. **D.** Violin plots representing the cell number in the meristem zone (*i.e.*, from the quiescent center to the first cell of the transition zone) at 5 DAG of WT, *ino80, een*, and *ino80 een* roots. Different letters denote significantly different groups (p < 0.05, *One*-way ANOVA with post-hoc Tukey HSD (Honestly Significant Difference) Test). **E.** Violin plots representing the length of root meristem (in µm) at 5 DAG of WT, *ino80, een*, and *ino80 een* roots. Different letters denote significantly different groups (p < 0.05, ANOVA with Tukey correction).

**Supplemental Figure 2:**
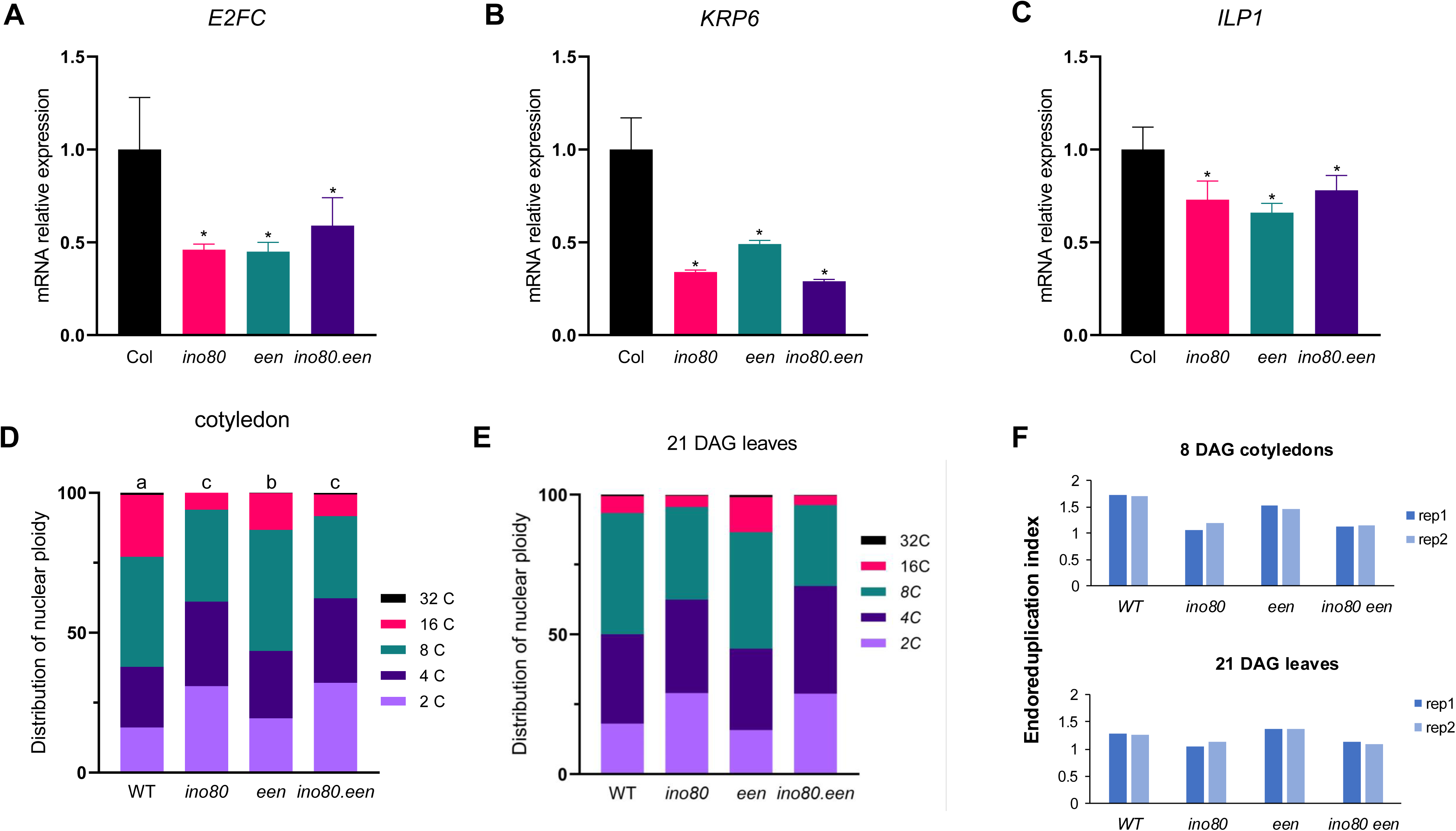
Ploidy level in WT, *ino80, een* and *ino80 een* plants. **A.** Histogram representing the mRNA steady state level (relative to WT plants) of *E2FC* in *ino80, een* and *ino80 een* Arabidopsis plants at 21 DAG. **B.** Same as **A.** for *KRP*. **C.** Same as **A.** for *ILP1*. (*p < 0.05; t-test). **D.** Histogram representing the distribution of 2C, 4C, 8C, 16C, or 32C nuclei from cotyledons of WT, *ino80, een* and *ino80 een* Arabidopsis cotyledons at 8 DAG. Different letters above columns indicate significant differences among different genotypes. (p < 0.05; Chi-squared test using WT as the expected frequency). **E.** same as **D.** from leaves 1 and 2 of WT, *ino80, een* and *ino80 een* Arabidopsis plants at 21 DAG. **F.** Histograms representing the endoreduplication index of WT, *ino80, een* and *ino80 een* Arabidopsis cotyledons at 8 DAG (upper panel) and in leaves 1 and 2 at 21 DAG (lower panel). Two independent biological replicates (rep1 and 2) are shown.

**Supplemental figure 3:**
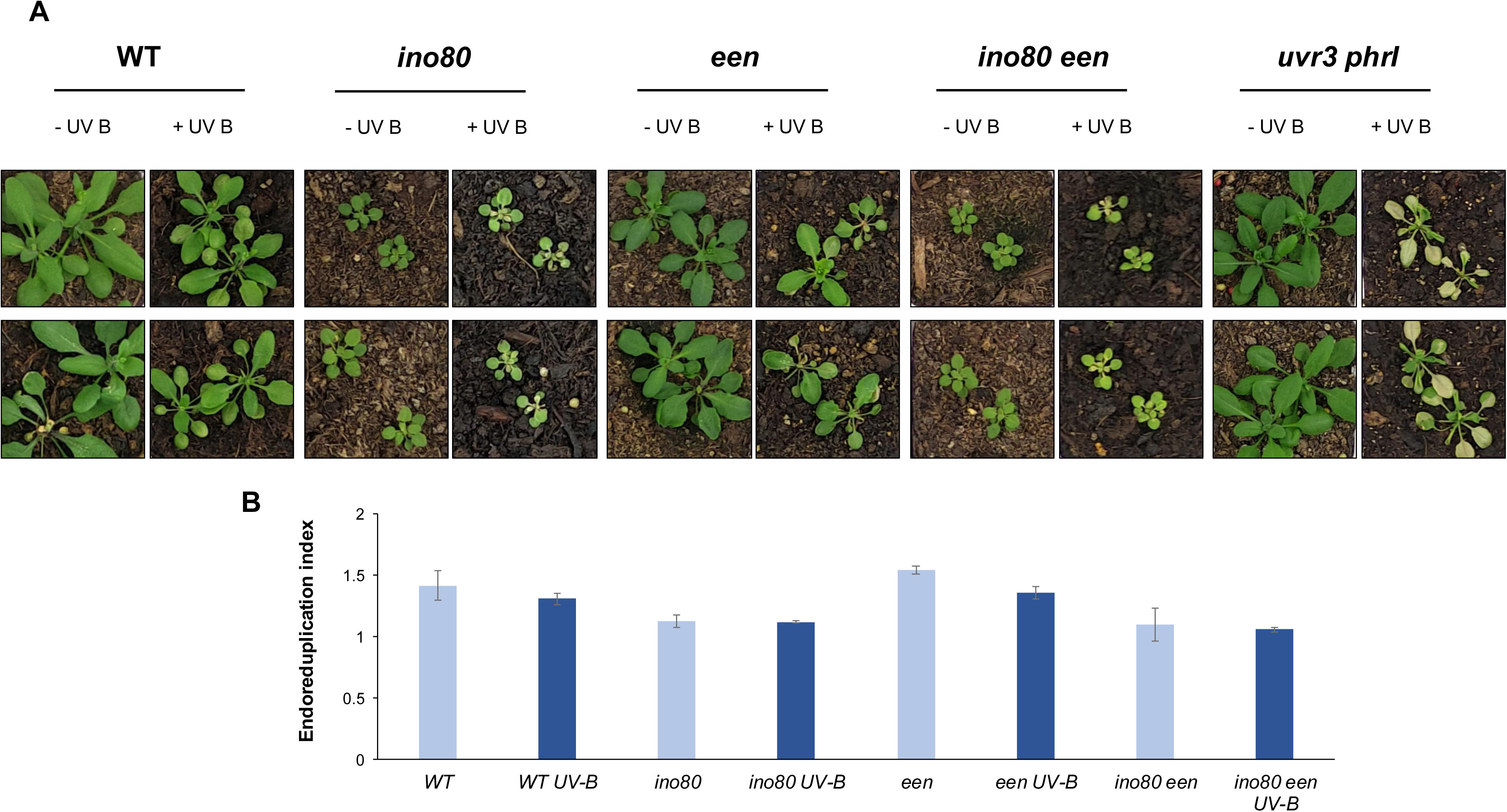
Phenotype of UV-B-treated WT, *ino80, een*, and *ino80 een* Arabidopsis plants. **A.** Rosette phenotype of UV-B-treated WT, *ino80, een*, *ino80 een* and *uvr3 phrI* Arabidopsis plants. **B.** Endoreduplication index of WT, *een*, *ino80* and *ino80 een* Arabidopsis plants ± UV-B (*p < 0.05; t-test).

**Supplemental Table 1.**
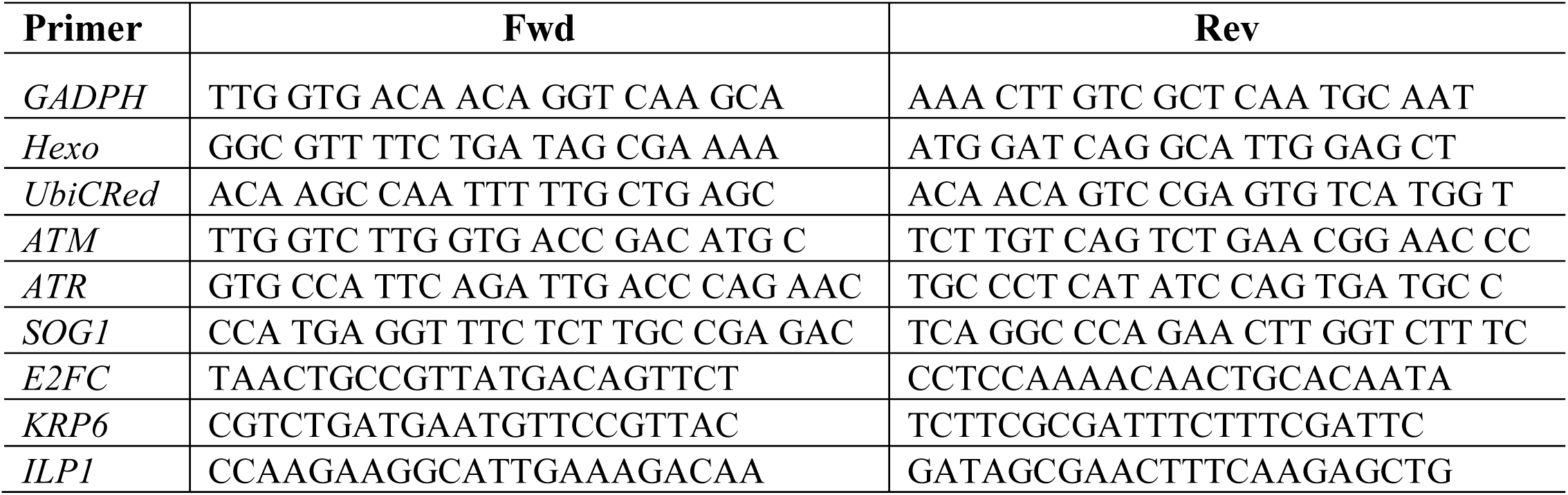
List of primers for qPCR.

**Supplemental Table 2.**
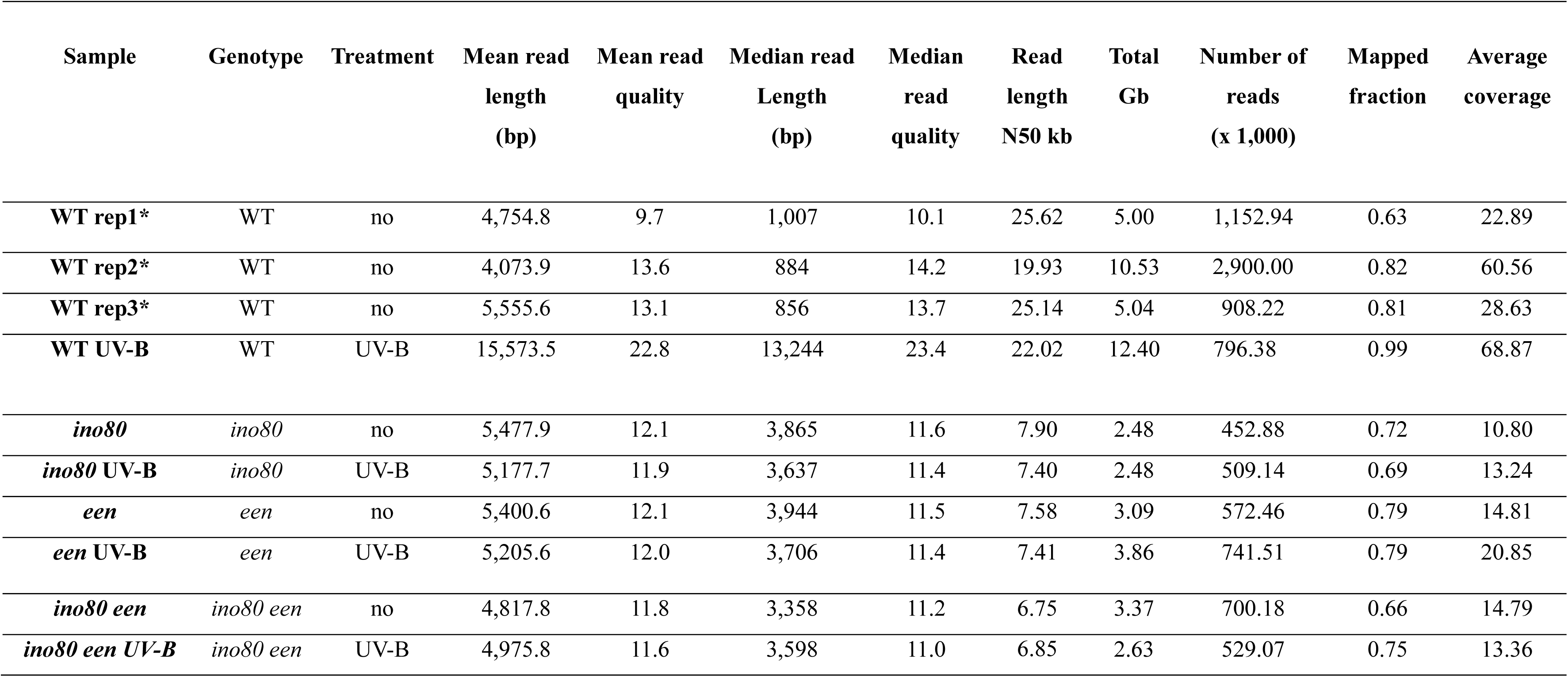
Sequencing statistics. *Sall et al. (2025)

